# Clock genes *period* and *timeless* control synaptogenesis in *Drosophila* motor terminals

**DOI:** 10.1101/2024.11.15.621522

**Authors:** María José Ferreiro, Soledad Astrada, Rafael Cantera, Santiago Ruiz

## Abstract

Some neurons undergo rhythmic morphological changes that persist in constant darkness and require the expression of clock genes. Flight motoneuron MN5, one of the best studied neurons of *Drosophila melanogaster* exhibits several examples of this type of circadian structural plasticity. During the morning, when the fly is active, synaptic boutons are larger than during the night when the fly is resting though more synaptic boutons and synapses are observed. Here, by comparing bouton numbers at different timepoints in normal flies and in flies carrying loss-of-function mutations in clock genes *timeless* (*tim*) or *period* (*per*), we investigate whether the rhythmic changes in numbers of boutons and synapses require the expression of these genes. Absence of *tim* expression abolished the rhythm in bouton number whereas absence of *per* expression appeared to increase the rhythm’s amplitude. This indicates that in normal flies TIM protein is necessary to drive the normal rhythm of bouton number and PER probably has a damping effect on it. In addition, it appears that *tim* and *per* expression normally act as inhibitor of synaptogenesis because their loss-of-function mutations caused over-proliferation of synapses. Unexpectedly, TIM and PER were expressed in different cells. TIM was found in the glial sheath wrapping the motoneuron’s axon and PER was predominantly found along the axon, suggesting that the control of the rhythmic change in bouton and synapse numbers requires interactions between different cell types.

## INTRODUCTION

The term “structural circadian plasticity” (Muraro et al., 2013) refers to a particular type of neuronal plasticity in which a neuron’s morphology changes in a predictable way, with a daily rhythmicity and under the control of the biological clock. It is a wide-spread phenomenon among many animal species and includes changes in axonal diameter, axonal branching, synapse numbers, synaptic vesicle size and distribution, density and connectivity between different pre- and post-synaptic partners (Pyza and Górska-Andrzejak, 2004; Mehnert and Cantera, 2011; Muraro et al., 2013; Frank and Cantera, 2014; Krezptowski et al., 2018; Frank, 2020). Among the proteins reported to participate in the control of circadian structural plasticity, two of the most studied are those encoded by the “clock genes” *timeless* (*tim*) and *period* (*per*) (Muraro et al., 2013; Krzeptowski et al., 2018; Cai & Chiu JC, 2022). TIM and PER proteins are fundamental components of the molecular mechanism that drives the clock, acting in auto-regulatory transcriptional loops, but they also exert other functions (Bell-Pedersen et al., 2005; Peschel and Helfrich-Förster, 2011; Jarabo and Martin, 2017; Pureumet al., 2019; Vipat and Moiseeva, 2024).

The innervation of flight muscles in *Drosophila melanogaster* provides a remarkable example of circadian neuronal plasticity. Motoneuron 5 (MN5), which innervates two of the very large longitudinal flight muscles, shows a daily change in size of synaptic boutons during the phase of rest/sleep (Mehnert et al., 2007). Measurement of bouton size in wild-type flies (*wt*), every three hours during a complete Light:Darkness (LD) cycle, showed a rhythmic increase in bouton size with a single peak in the morning when the fly increment its locomotor activity (Mehnert and Cantera, 2008).

Three lines of evidence indicate that the rhythmic enlargement of bouton size is not a simple consequence of rhythmic changes in synaptic activity. Firstly, the morning enlargement does not happen again later in the day, when the fly has a second daily “bout” of locomotion activity (Wheeler et al., 1993; Helfrich-Forster, 2005); secondly, it persists in *shibire* mutant flies whose synaptic activity is experimentally eliminated and lastly, it persists during 48 hours in wild-type flies that are rendered motionless by decapitation (Mehnert and Cantera, 2008).

The MN5 axonal terminal also shows a daily change in the number of boutons and synapses with the largest number of synapses reported at midnight (Ruiz et al., 2013), when the fly does not fly. A synaptic bouton can have one or several synapses, but it can also lack synapses (referred as “empty boutons” in Ruiz et al., 2013). Normally, half of the synaptic boutons of the MN5 are “empty boutons” at midnight and this proportion is reduced at midday (Ruiz et al., 2013).

The main biological clock resides in the brain (Helfrich-Förster, 2005; Roenneberg and Merrow, 2016; King and Sehgal, 2018). However, the rhythmic change of MN5 bouton size continues in decapitated flies (Mehnert and Cantera, 2008), indicating that it can be driven from one or several peripheral pacemakers, as it is the case for other circadian rhythms (Giebultowicz, 2001; Brown et al., 2019). Among other tissues and cells with pacemaker properties that could influence circadian structural plasticity of MN5, glia cells are interesting due to their participation in circadian rhythms of neuronal morphology and synapse plasticity (Gorska-Andrzejak et al, 2013; Herrero et al., 2017; Chi-Castañeda and Ortega, 2018; McCauley et al., 2020; Gamulewicz et al., 2022).

It was still not known whether the rhythmic change in the number of boutons and synapses also requires the expression of clock genes, as it is the case for the rhythmic change in bouton size. In the present work, we quantify the numbers of MN5 boutons and synapses at different time points of the day in flies carrying null mutations in *tim* or *per* kept in constant darkness. We found that the rhythmic increment in the number of boutons that normally occurs during the subjective night, persisted in *per* mutants but was abolished in *tim* mutants. Both mutations incremented the number of boutons with synapses, and particularly the proportion of boutons with two or more synapses.

## MATERIALS AND METHODS

### Fly strains and experimental conditions

The fly strains of *Drosophila melanogaster* used here were *Oregon R* (*wt*), the clock null mutants *tim*^*01*^ and *per*^*01*^ and the transgenic lines *repo-gal4* (a specific driver for glial cells, Xiong et al., 1994; Halter et al., 1995) and *uas-cd8-gfp* (Lee and Luo, 1999). Flies from the *repo-gal4* strain were crossed with *uas-cd8-gfp* flies to label glial cells with specific, strong and membrane-bound green fluorescence (GFP).

All strains were maintained on standard *Drosophila* medium at 25°C and kept at LD cycles of 12:12 hours for at least three days before being transferred to Darkness:Darkness (DD) conditions. For DD experiments, the light was switched off at Zeitgeber time 0 (ZT0) the first day of the experiment setting the circadian time 0 (CT0). All experiments were done using 4-5-days old female flies since they exhibit strong rhythmicity in the size of boutons (Mehnert et al., 2007; Mehnert and Cantera, 2008) and in the number of synaptic vesicles (Ruiz et al., 2010). Two time points were studied under DD conditions: circadian time 19 (CT19) corresponding to subjective midnight (middle of the sleep phase) and circadian time 7 (CT7) corresponding to subjective midday. The flies were anesthetized with nitric oxide (Sleeper TAS, INJECT+MATIC, Switzerland), decapitated with a sharp needle and thereafter the dorsal portion of the thorax was dissected and fixed as described below. The number of flies of each genotype analyzed in each condition is expressed in Figs 2 and 3 captions.

**Figure 1.**
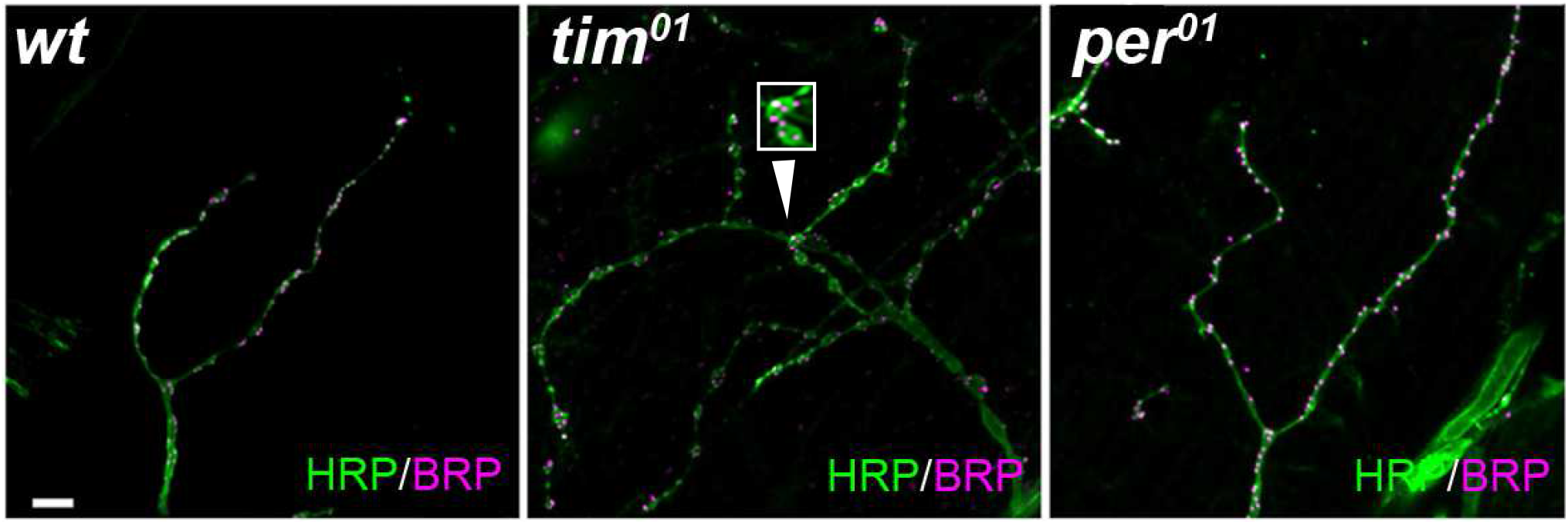
Neuroanatomy of MN5’s motor terminals in *timeless* and *period* mutant flies. The neuronal membrane was stained with anti-HRP (green) and the synapses with anti-BRP (magenta). Only a small portion of each MN5 terminal is shown here (an entire MN5 axonal terminal has hundreds of branches) but the images were selected to be representative of each phenotype and the quantitative method used here gives consistent results for each genotype (Ruiz et al., 2013). Scale bar= 5μm.

**Figure 2.**
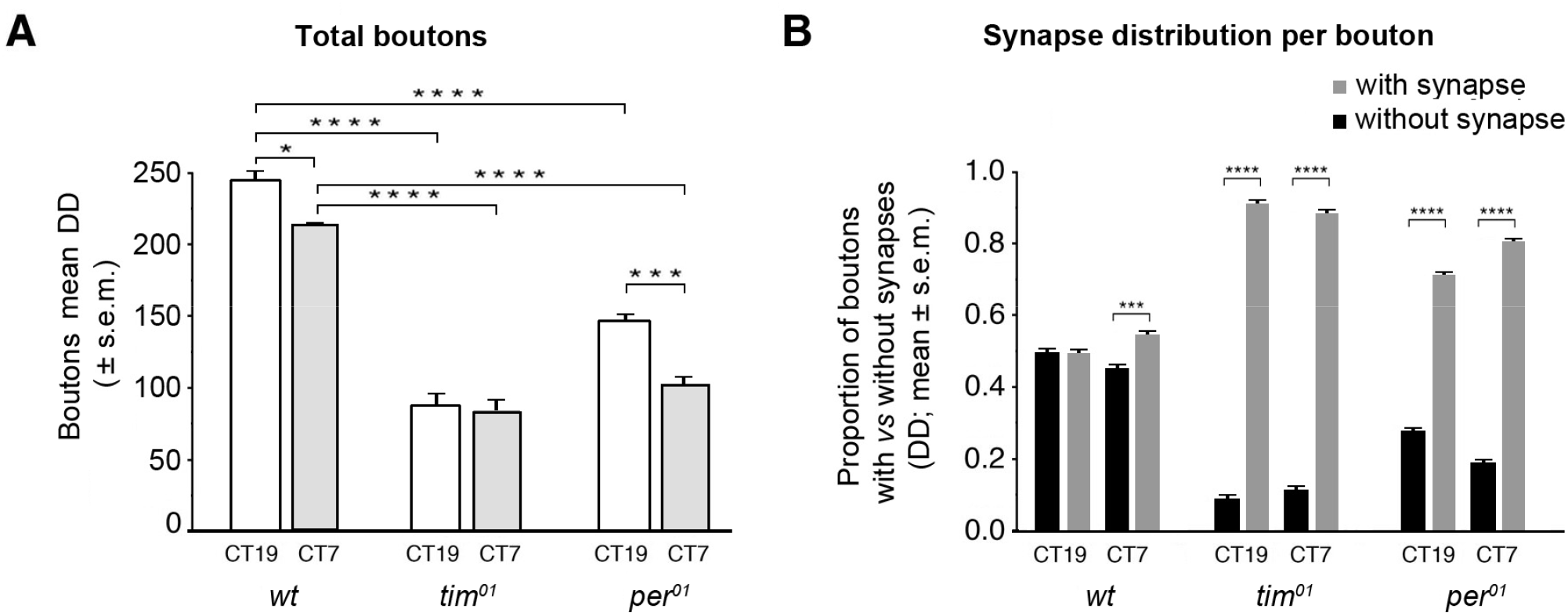
Null mutations in the genes *timeless* and *period* interfere with the daily rhythm in the number of synaptic boutons and synapses. (A) The graph represents the total number of boutons in the MN5 motor terminal of *Drosophila* (mean ± SEM) measured either at subjective midday (CT7) or subjective midnight (CT19) in *wt* or in *tim*^*01*^ and *per*^*01*^ clock mutant flies reared in DD. Normally *wt* flies have more boutons at CT19 than CT7 (CT19 *vs* CT7= 240 ± 10.5 *vs* 212± 8.8, Student *t* test, P= 0.04, n= 33 and 37 flies per time point, respectively). This change was absent in *tim*^*01*^ mutants (CT19 *vs* CT7= 88 ± 6.09 *vs* 85 ± 6.09, Student *t* test, n.s., n= 14 and 16 flies per time point, respectively) but was even larger in *per*^*01*^ mutants (CT19 *vs* CT7= 146 ± 8.3 *vs* 104 ± 6.7, Student *t* test, P= 0.0007). Both mutants had fewer boutons than *wt* flies. (B) Graphic representation of the proportion of boutons with synapses *vs* without synapses (mean ± SEM, relative to the total number of boutons found per time point in each experiment), in *wt, tim*^*01*^ and *per*^*01*^ mutant flies. In *wt* flies, the proportion of boutons both with and without synapses was 50:50 (with:without) at subjective midnight (CT19: 0.50 ± 0.01 *vs* 0.50 ± 0.01, respectively, Student *t* test, n.s.) and 55:45 (with:without) at subjective midday (CT7: 0.55 ± 0.01 *vs* 0.45 ± 0.01, respectively, Student *t* test, P= 0.00012). The two clock mutants had an abnormally high proportion of boutons with synapses regardless the moment of the day. The highest proportion was found in *tim*^*01*^ mutants (CT19: 0.91 ± 0.02 *vs* 0.09 ± 0.02, Mann-Whitney U test, P= 0.000007, CT7: 0.89 ± 0.01 *vs* 0.11 ± 0.01, Mann-Whitney U test, P= 0.000002), followed by *per*^*01*^ mutants (CT19: 0.72 ± 0.02 *vs* 0.28 ± 0.02, Mann-Whitney U test, P= 0.000000, CT7: 0.81 ± 0.02 *vs* 0.19 ± 0.02, Mann-Whitney U test, P= 0.000003) (*) p< 0.05; (***) p< 0.001; (****) p< 0.0001; n.s.= non-significant.

**Figure 3.**
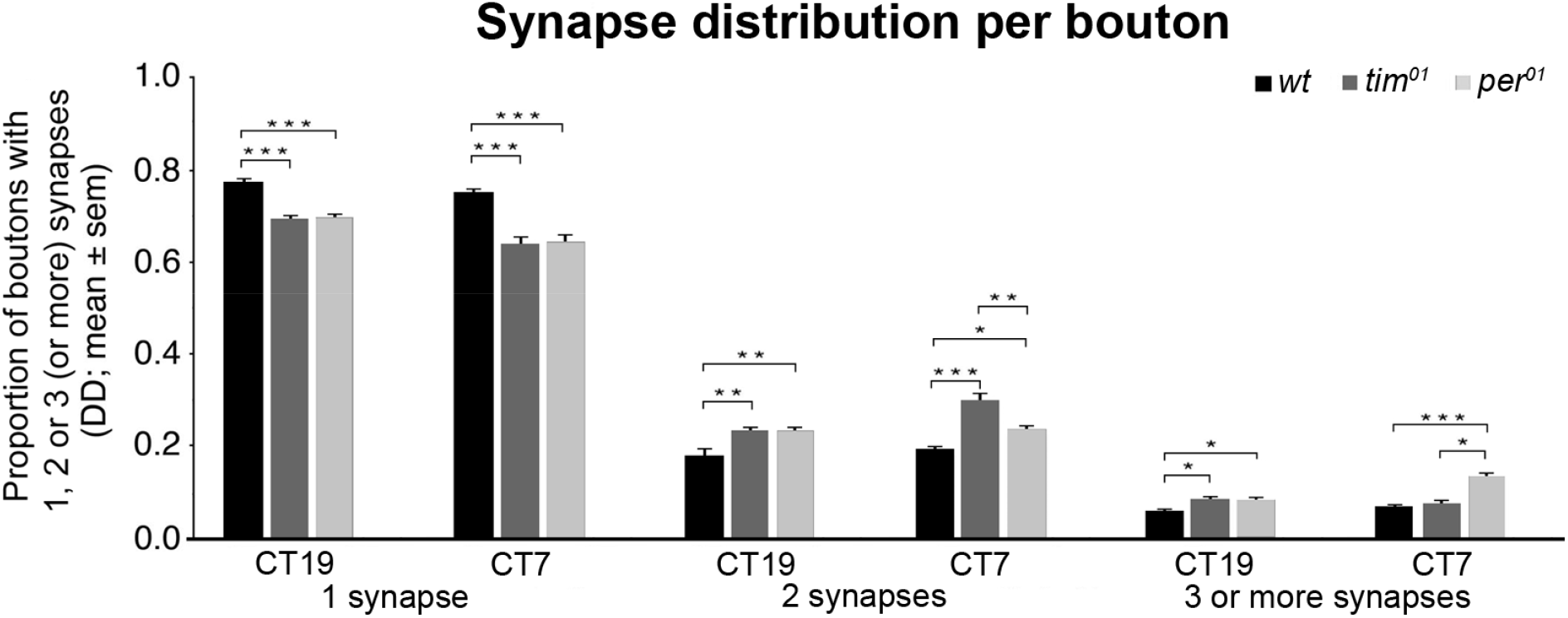
Synapse distribution among MN5 boutons in *Drosophila* mutants *timeless* or *period*. The graph represents the proportion of MN5 boutons with one, two or three or more synapses (mean ± SEM), counted either at subjective midday (CT7) or subjective midnight (CT19) in *wt* or in clock mutant flies *tim*^*01*^ and *per*^*01*^ reared in DD. In all genotypes the great majority of boutons with synapses had only one synapse, few had two and even fewer had three or more synapses. In *wt* flies the relative proportion of these three categories was 75:20:5 (Kruskal-Wallis test, CT19: H(2,N= 99)= 85.47, P= 0.000; CT7: H(2,N= 111)= 96.79, P= 0.000). In both clock mutants, on the other hand, the proportion of boutons with more than one synapse increased, shifting to approximately 70:20:10 at CT19, (the same as for *wt* flies kept in LD, see Ruiz et al., 2013). At CT7, the proportion shifted to approximately 65:30:5 in *tim*^*01*^ mutants (Kruskal-Wallis test, CT7: H(2,N= 48)= 41.63, P= 0.000) and to approximately 65:20:15 in *per*^*01*^ mutants (Kruskal-Wallis test, CT7: H(2,N= 45)= 36.12, P= 0.000). (*) p< 0.05; (**) p< 0.01; (***) p< 0.001; (****) p< 0.0001.

### Immunohistochemistry and microscopy

To count the total number of boutons the samples were doubly stained with a rabbit polyclonal anti-Horseradish Peroxidase serum which labels motoneuron plasma membrane (by recognizing the glycoprotein 3-alpha-L-fucosyltransferase; Jan and Jan, 1982; Sun and Salvaterra, 1995) and mouse monoclonal antibody nc82 (Hofbauer, 1991) which recognizes the protein Bruchpilot (BRP, an ELK/CAST homologue), which is an important component of the T-bar, a presynaptic component of insect synapses (Trujillo-Cenóz, 1969; Prokop and Meinertzhagen, 2006; Wagh et al., 2006; Kittel et al., 2006).

The thorax was opened along the mid-sagittal plane in a droplet of phosphate-buffered saline (PBS, 0.1M at pH 7.4). Only one hemithorax per fly was processed. Samples were fixed on ice-cold 4% (w/v) paraformaldehyde in PBS for 2.5 hours, tissues were then permeabilized using 0.3% Triton X100 in PBS (PBST) and unspecific binding sites were blocked with 1% Normal Goat Serum in PBST 0.5% bovine serum albumin. Samples were then incubated with anti-HRP (1:600, Jackson ImmunoResearch, Cat# 323-001-021, RRID: AB_2340254) and anti-BRP (1:100, DSHB, Cat# nc82, RRID: AB_2314866) antibodies over-night at room temperature (RT). The next day, samples were washed 4×15 minutes in PBST and then incubated with goat anti-rabbit secondary antibodies conjugated to Alexa 488 (1:1000, Cat# A32731, RRID AB_2633280, Thermo Fisher Scientific) and goat anti-mouse secondary antibodies conjugated to Cy3 (1:1000, Cat# 115-001-001, RRID: AB2338442, Jackson ImmunoReseach, Inc.) for 2 hours at RT. Samples were then washed 4×15 minutes in PBST and 2×10 minutes in PBS. Finally, the indirect flight muscles 5 (IFM5) and 6 (IFM6), which are jointly innervated by MN5, were mounted in 80% glycerol PBS. Double-staining assays of the neuromuscular junction (NMJ) of the MN5 with rabbit and rat polyclonal antiserum specific for TIM (UP991 1:1000 and UPR41 1:100, provided by Amita Sehgal) and rabbit polyclonal antiserum specific for PER (1:700, provided by Ralf Stanewsky), were performed following the same general protocol previously described, to detect TIM and PER proteins respectively.

### Image acquisition and quantification of boutons and synapses

Samples were examined using a laser confocal microscope (LCM) Olympus FV300. Two images per fly were taken for bouton and synapse blind quantification. Images analysed were a result of a maximum intensity Z projection of optical sections scanned at intervals of 0.3 μm with a 60x lens and a digital zoom of 3.5 and analyzed with ImageJ.

For each sample, the total number of boutons and of boutons with or without synapses was counted “in blind” by the same person to avoid personal bias. The resulting numbers are only a fraction of the total number of synapses in each sample (See below).

### Statistical analysis

Student *t*-tests were performed when assumptions for parametric test were accomplished (normality using Shapiro-Wilk W test and homoscedasticity using Levenés test). If these assumptions were not achieved, non-parametric Kruskal-Wallis and Mann-Whitney U tests were performed instead. Statistical significance was set at 0.05, 0.01,0.001 and 0.0001. Analyses were done using Statistica 12 software (Statsoft).

## RESULTS

### General neuroanatomy of MN5’s motor terminals in timeless and period mutant flies

The axonal terminal of the MN5 extends over two very large muscles and penetrate them deeply with hundreds of terminal branches that carry a total of approximately 4000 boutons containing 3000 BRP-positive synapses (Ruiz et al., 2013). It is therefore not feasible to count the total number of boutons or synapses in this axonal terminal, but we have developed a quantification method based on small, uniform and representative microvolumes of flight muscle that produces results with excellent replicability and reproducibility (Ruiz et al., 2013).

The monoclonal antibody nc82 that we used to label Bruchpilot-positive synapses, binds exclusively to the presynaptic site of synapses as shown with immunogold staining for electron microscopy (Fouquet et al., 2009; Hamanaka and Meinertzhagen, 2010). Anti-BRP staining appears as strong fluorescent “spots” in a practically negative background and has been widely used in *Drosophila* to assess the number of synapses in the NMJ of different muscles (Wagh et al., 2006; Ataman et al., 2008; Fouquet et al, 2009; Ruiz et al., 2013).

The proportion of boutons per time point was calculated as the number of boutons found in each time-point divided by the number of flies. The proportion of boutons with vs. without synapses was also calculated per time point, relative to the total number of boutons found at each time point. Boutons with synapses were classified into three classes: boutons with one synapse, boutons with two synapses and boutons with three or more synapses. Thereafter the percentage of each class was calculated. All parameters were graphically represented as the mean ± SEM.

Previous studies reported that the terminal of the MN5 suffers over-branching and abnormal axonal morphology in *tim* mutants but reduced number of branches in *per* mutants (Mehnert et al., 2007). Similar phenotypes were reported for clock neurons in the brain of adult flies (Fernández et al., 2008). Consistent with these reports, here we found that *tim*^*01*^ mutants motor terminals showed over-branching and larger boutons compared to that of *wt* flies, whereas in *per*^*01*^ mutants the terminals had reduced branching and smaller boutons compared to *wt* flies (Fig.1). We also found sporadic cases of “sprouting” in the MN5 terminals of *tim* mutant (data not shown).

### The rhythmic change in the number of synaptic boutons has larger amplitude in per mutants but is abolished in tim mutants

The NMJ made by the MN5 on its two target muscles has normally more boutons at midnight compared to midday both in LD and DD conditions (Ruiz et al., 2013). This feature was confirmed here in the DD samples of *wt* flies studied when measured during two consecutive days (CT19 *vs* CT7 Fig. 2A). The amplitude of this change was significantly larger in *per* mutants but absent in *tim* mutants (Fig. 2A).

It has been reported that at midnight half of the boutons of the MN5 axonal terminal contain no synapses and that the proportion of boutons with synapses increases during the day in LD or the subjective day in DD (Ruiz et al., 2013). Here we confirmed these features in *wt* flies kept in DD (Fig.2B). A large increment in the proportion of boutons with synapses was observed in *tim* and *per* mutants at both timepoints, reaching approximately 90% in *tim*^*01*^ mutants and 80% in *per*^*01*^ mutants during the subjective midnight (Fig.2B).

In summary, TIM appears to be necessary for the rhythmic increment in bouton numbers during the night whereas PER appears to have a damping effect on the rhythm’s amplitude. More interestingly, it appears that both proteins normally exert an inhibitory function on synapse formation because their absence results in a greater proportion of boutons carrying synapses.

### Timeless and Period control the distribution of synapses among boutons

Normally at midnight, only half of the boutons found along MN5 terminals of *wt* flies kept in DD conditions have synapses (Fig.2B). Among these boutons, the relative proportions of boutons with either one, two or more synapses is 75:20:5 (Ruiz et al., 2013). Here we found a significant shift on this distribution in samples from clock mutant flies, with a reduction in the proportion of boutons with one synapse and an increment in the proportion of boutons with two, three or more synapses (70:20:10; Fig.3). These proportions were like those reported for *wt* flies kept in LD (Ruiz et al., 2013). At CT7 the proportion was approximately 65:30:5 in *tim*^*01*^ mutants and approximately 65:20:15 in *per*^*01*^ mutants.

### Timeless and Period are differentially expressed in the neuromuscular junction of the MN5

The observation that loss-of-function mutations in either *tim* or *per* have different effects on MN5 synapses lead us to investigate the possibility of differential spatial expression of the corresponding proteins. To this end, the localization of TIM and PER proteins in the NMJ of the MN5 was investigated with laser confocal microscopy using well characterized antibodies in transgenic flies expressing GFP in glial cells. It was found that PER was predominantly located along the axon and its branches, whereas TIM was predominantly localized in the glia that surrounds the axon and its branches (Fig. 4).

**Figure 4.**
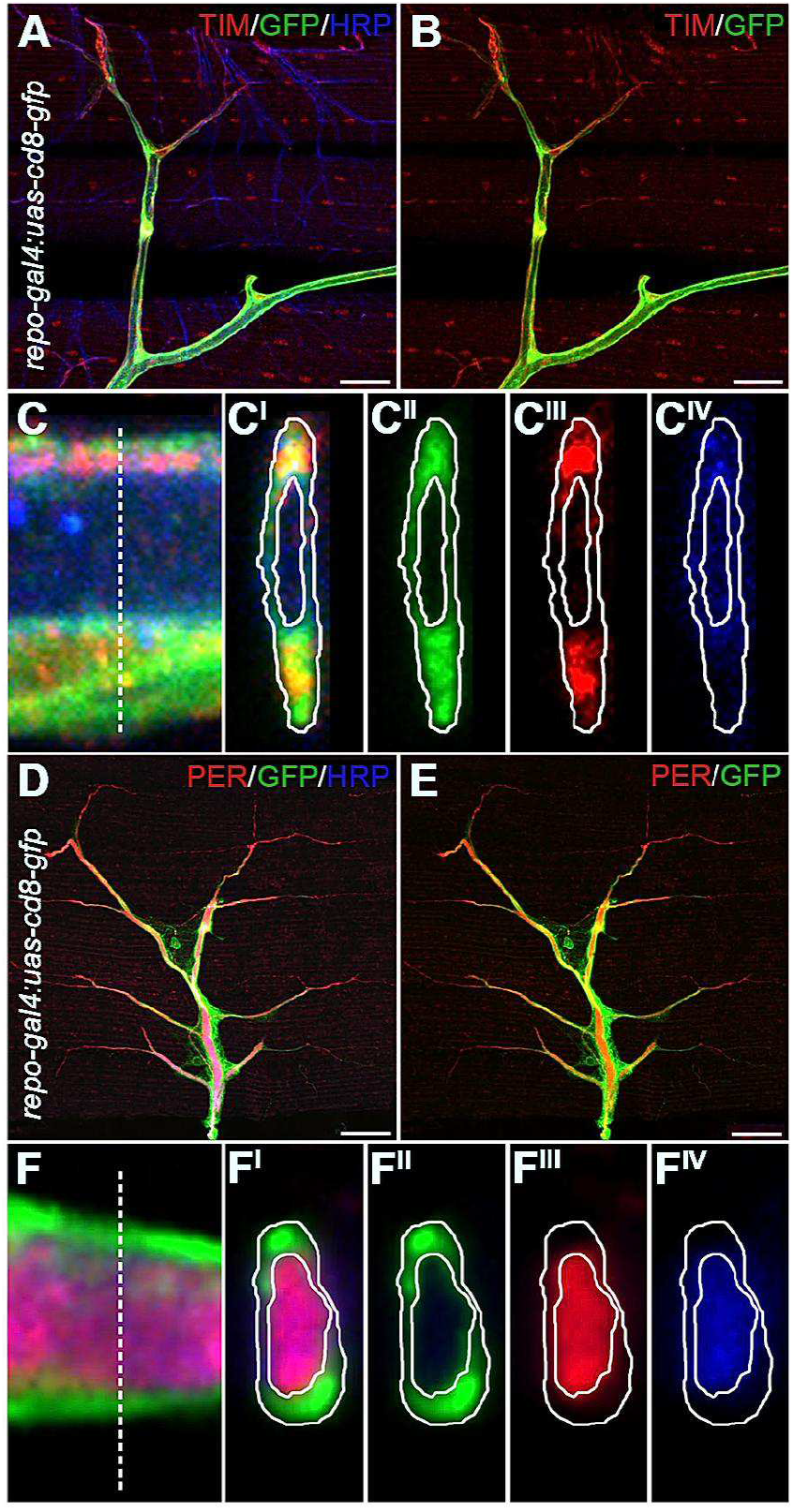
Clock proteins TIM and PER are differentially expressed in different cellular components of the *Drosophila* MN5 NMJ. All images show immunostainings of flight muscle tissue expressing membrane-bound GFP in all glia (*repo-gal4*; *uas-cd8-gfp*) associated with the MN5’s motor terminal, corresponding to CT19 samples. (A-C) TIM was expressed preponderantly in the glia wrapping the axon. (A) Representative view of the NMJ immunostained for TIM (anti-TIM, Red) and for a neuronal membrane protein (anti-HRP, Blue). Glia cells were identified by their transgenic expression of green fluorescent protein anchored to the membrane through the CD8 protein (GFP, Green). (B) Same view as in (A) after elimination of the blue channel for clarity. (C) Single longitudinal focal plane of an orthogonal reconstruction of a series of confocal sections taken at small intervals along the Z-axis to show the localization of the three markers simultaneously (green= glia; red= TIM; blue= MN5), (C^I^): orthogonal view of the three markers altogether and orthogonal views of the three markers separately: (C^II^= glia), (C^III^= TIM) and (C^IV^= MN5). The dashed line in (C) indicates the plane used for the orthogonal reconstructions. (D-F) The main expression of PER was found in the axon of the MN5. (D) Representative view of the MN5 NMJ immunostained for PER (anti-PER, Red) and for a neuronal membrane protein (anti-HRP, Blue). Glial cells express specifically green fluorescent protein anchored to the membrane through the CD8 protein (GFP, Green). (E) Same view as in (D) after elimination of the blue channel for clarity. (F) Single longitudinal focal plane of an orthogonal reconstruction of a series of confocal sections taken at small intervals along the Z-axis to show the localization of the three markers simultaneously (green= glia; red= PER; blue= MN5), (F^I^): orthogonal view of the three markers altogether and orthogonal views of the three markers separately: (F^II^= glia), (F^III^= PER) and (F^IV^= MN5). The dashed line in (E) indicates the plane used for the orthogonal reconstructions (Scale bar = 20μm).

## DISCUSSION

The experimental evidence presented here indicates that the clock genes *tim* and *per* participate in the control of a circadian rhythm of structural plasticity of the terminal that motorneuron MN5 forms on flight muscles in *Drosophila*. This rhythm normally includes a daily change in the number of synaptic boutons and in the number and distribution of synapses (active sites) (Fig. 5).

**Figure 5.**
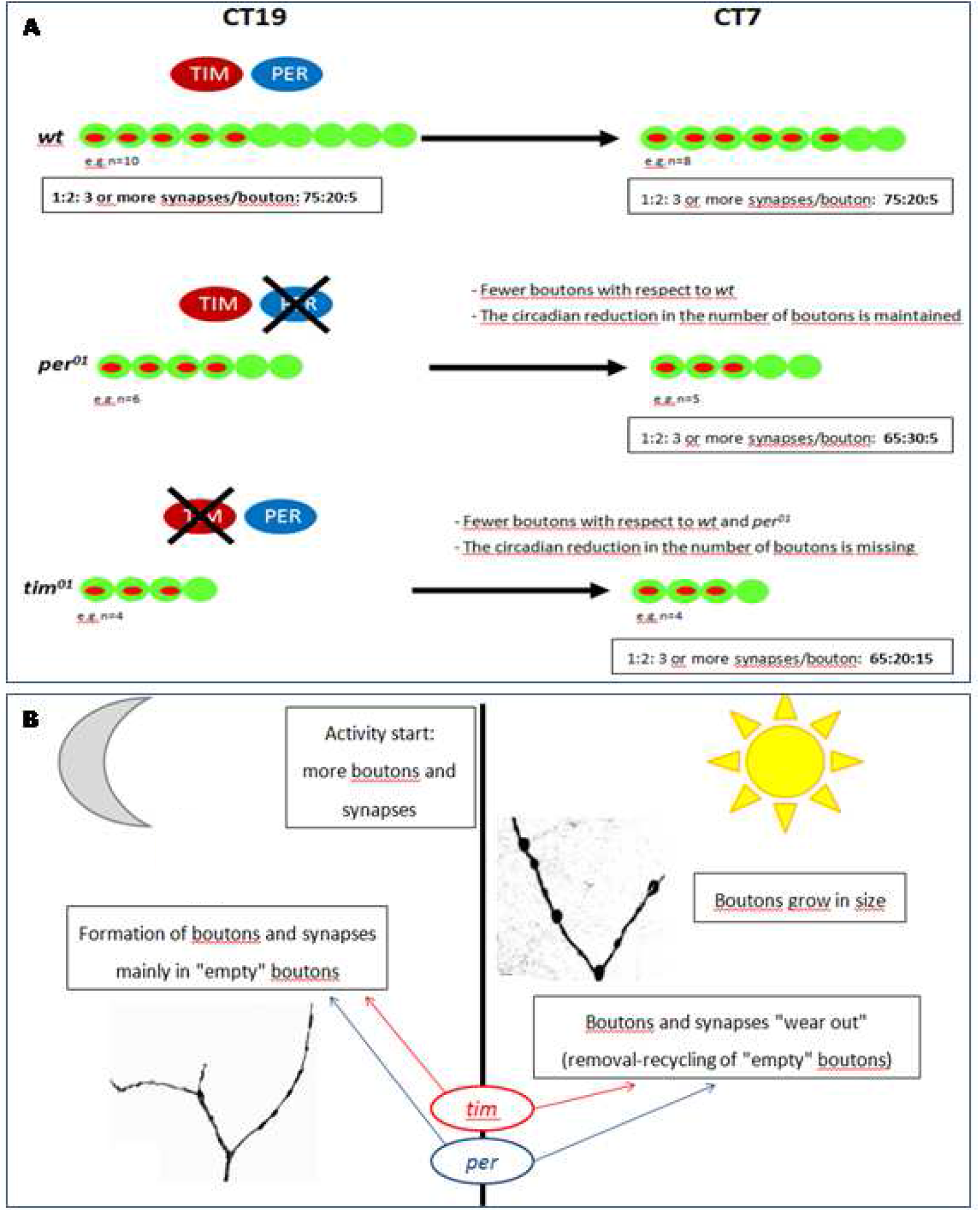
Circadian plasticity of MN5 synapses. A) Absence of *tim* expression (*tim*^*01*^) abolish the rhythm in bouton number, whereas absence of *per* expression (*per*^*01*^) seems to increment the rhythm’s amplitude compared to *wt*. TIM and PER will normally act as inhibitors of synaptogenesis, because their absence causes over-proliferation of synapses. B) Towards the end of the night, when the fly will start its morning peak of activity, the MN5 will have a relatively large number of boutons with synapses, some of which were assembled during the night. During the day, as the fly flies, MN5 boutons will grow probably through the fusion of membrane from synaptic vesicles emptied during activity.

Our findings demonstrate that both TIM and PER are necessary for this daily reorganization of the MN5 terminal, and that each of the two protein is expressed in a different cell type (Fig. 4) suggesting that they participate in a mechanism that controls structural synaptic plasticity through interactions between glia and neurons. It appears that one of the normal functions of both proteins is to promote the formation of synaptic boutons and inhibit the formation of synapses (Figs. 2 and 3).

During the night, TIM contributes to a mechanism that drives a rhythmic increment in the number of synaptic boutons relative to midday, and PER controls the amplitude of that rhythm. Absence of *tim* expression abolish the rhythm in bouton number, whereas absence of *per* expression results in an increment the rhythm’s amplitude compared to *wt* (Fig.5A).

MN5 terminal of flies maintained in DD normally comprises equal proportions of boutons with or without synapses at subjective midnight but larger proportion of boutons with synapses at subjective midday (Ruiz et al., 2013 and this study). The present study suggest that this rhythmic increment of boutons carrying synapses during the day is normally repressed by TIM and PER. It appears that TIM and PER normally act as inhibitors of synaptogenesis, because the loss-of-function mutations in their genes cause over-proliferation of synapses (Fig. 5A).

A striking result regarding *per* mutant flies is that they showed a rhythmic change in the number of boutons under DD conditions. Such result contradicts previous observations for the circadian rhythms in locomotor activity, neuronal morphology, size in synaptic boutons, and other circadian rhythms which are either abolished or seriously diminished when these mutants are kept in DD conditions (Konopka & Benzer, 1971; Sakai & Ishida, 2001; Cascallares et al., 2018). However, a small percentage of *per* mutants still exhibit DD rhytmicity under certain conditions (Schlichting et al., 2019). For example, Nippe (2015) reported that under DD about 19 % of *per* null mutants have a detectable rhythm in mean visual contrast sensitivity in the visual centers. There are also reports of biological rhythms without *per* transcription (Millius et al., 2019).

It was already known that loss-of-function in *tim* and *per* genes caused different mutant phenotypes in the MN5 (Mehnert et al., 2007) as well as in other neurons (Fernandez et al., 2008; Gorostiza et al., 2014; Herrero et al., 2017). Our findings add to these previous results and challenge the view that *per* and *tim* control structural circadian plasticity exclusively through their canonical role as molecular components of the biological clock (i. e. acting as heterodimers that regulate transcription of the same target genes).

A finding that reinforces the probability of an alternative function is the differential localization of TIM and PER proteins. Both proteins are localized in different cell types, neuron and glia, suggesting that the mechanism controlling the daily reorganization of synapses involves interactions between at least two cell types.

The initial hypothesis proposed that the rhythmic reorganization of MN5 terminals was controlled by a clock-gene dependent, non-autonomous mechanism probably driven from a peripheral pacemaker located in the MN5 itself, glia, its target muscles or interneurons (Mehnert et al., 2007; Mehnert and Cantera, 2008).

Whereas the participation of muscle or interneurons has yet to be investigated, our results suggest that the control of the daily reorganization of the MN5 synaptic terminal involves an interaction between the neuron and its associated glia.

In general, it is firmly established that glial cells influence neuronal morphology and synapse formation in *Drosophila* as in other animal species (Bialas and Stevens, 2012; Fuentes-Medel et al., 2012; Kerr et al., 2014) and control a variety of circadian rhythms (Jackson, et al., 2015; You et al., 2018) including the circadian remodeling of neurons (Herrero et al., 2017). Particularly, in the MN5, the occurrence of an interaction between the neuron and its associated glia is supported by previous findings where the MN5 axon is shown to be tightly wrapped by concentric layers of glia. Thinner axonal branches located deep inside the flight muscles are also surrounded by glia (Ruiz et al., 2013). Cell-type specific knock-down of *per* or *tim* in each of the relevant cell types could be used to test this possibility.

Previously, we proposed that the change in the number of boutons and synapses of the MN5 between night and day could reflect a daily cycle of synapse recycling. It was suggested that during the day, when flight muscles are in use, synapses are “wasted” and disassembled, followed by a phase of synapse assembly during the night, when the fly is at rest (Ruiz et al., 2013). Taking into consideration the new data presented here, we propose a more detailed model (Fig. 5). Towards the end of the night, when the fly will start its morning peak of activity, the MN5 will have a relatively large number of boutons with synapses, some of which were assembled during the night. During the day, as the fly flies, MN5 boutons will grow probably through the fusion of membrane from synaptic vesicles emptied during activity. This may happen twice a day, because there is a tight correlation between the morning and the evening peaks of locomotion activity and the diameter and relative proportions of synaptic vesicle pools (Ruiz et al., 2010). Furthermore, during the wake phase some MN5 boutons and synapses will be damaged and dismantled (“wasted”) and/or recycled along a global program of ordered disassembly, elimination and recycling. This process will mainly affect the subpopulation of boutons without synapses if “wasted synapses” can be eliminated without eliminating the entire bouton (we envision that the population of boutons without synapses includes those which have yet not acquired their first synapse and those that have lost their synapses). In brain interneurons that are also known to reorganize their axonal terminals along a circadian cycle (Fernandez et al., 2008; Gorostiza et al., 2014) it is not the number of synapses that changes between day and night but their relative dispersion, with larger inter-synaptic distances during the morning (Hofbauer et al., 2024).

Most studies reporting the number of synapses (active sites with a T-bar and a cluster of synaptic vesicles in the presynaptic site) in the NMJ on different somatic muscles revealed that a single bouton bears tens of synapses (Chou et al., 2020; Südhof, 2021; Newman et al., 2022). It seems clear that the NMJ of the MN5 differs markedly from this pattern because the great majority of its boutons contain either none or only one synapse. The finding of an enlarged quote of synapse-bearing boutons in both mutants (Fig.2B) indicated that the absence of either protein (TIM or PER) caused a synaptogenic phenotype. The synaptogenic phenotype reported here represents a clear deviation of the normal pattern of synapse distribution towards more boutons with several synapses, suggesting that *tim* and *per* genes normally control synapse distribution among boutons, limiting the addition of synapses to “empty” boutons and specially the addition of a second or third synapse to boutons that already have one synapse.

## ACKNOWLEDGEMENTS

The nc82 antibody, developed by Dr. Buchner, was obtained from the Developmental Studies Hybridoma Bank developed under the auspices of the NICHD and maintained by The University of Iowa, Department of Biology, Iowa City, IA 52242. We express our gratitude to Dr. María Fernanda Ceriani for providing *tim* and *per* mutant strains, and Drs. Ralf Stanewsky and Amita Sehgal for sharing anti-PER and anti-TIM antibodies. RC and SR acknowledge funding from PEDECIBA, ANII, SNI and grant 058-54 from Programa de Desarrollo Tecnológico.

